# eIF2B extends lifespan through inhibition of the integrated stress response

**DOI:** 10.1101/2020.08.13.244970

**Authors:** Maxime Derisbourg, Laura Wester, Ruth Baddi, Martin S. Denzel

**Affiliations:** Max Planck Institute for Biology of Ageing, D-50931 Cologne, Germany; CECAD - Cluster of Excellence, University of Cologne, D-50931 Cologne, Germany; Center for Molecular Medicine Cologne (CMMC), University of Cologne, D-50931 Cologne, Germany

## Abstract

Protein homeostasis is modulated by stress response pathways and its deficiency is a hallmark of aging. The integrated stress response (ISR) is a conserved stress-signaling pathway that tunes mRNA translation via phosphorylation of the translation initiation factor eIF2. ISR activation and translation initiation are finely balanced by eIF2 kinases and by the eIF2 guanine nucleotide exchange factor eIF2B. However, the role of the ISR during aging remains unexplored. Using a genomic screen in *Caenorhabditis elegans*, we discovered a role of eIF2B and the eIF2 kinases in longevity. By limiting the ISR, these mutations enhanced protein homeostasis and increased lifespan. Consistently, full ISR inhibition using phosphorylation-defective eIF2α or pharmacological ISR inhibition prolonged lifespan. Lifespan extension through ISR inhibition occurred without changes in overall protein synthesis, and depended on enhanced translational efficiency of the kinase KIN-35. Evidently, lifespan is limited by the ISR and its inhibition may provide an intervention in aging.

---

Aging is defined as the progressive loss of physiological integrity accompanied by reduced cellular, organ, and systemic performance. It is characterized by cellular hallmarks such as stem cell exhaustion, genomic instability, deregulated nutrient sensing and loss of protein homeostasis^1^. Thus, aging is the main risk factor for neurodegenerative disorders, cancer and metabolic syndrome. The aging process can be modulated by environmental and genetic factors, and several evolutionary conserved biological processes have been implicated in lifespan regulation^2^. Failure of protein homeostasis is an early event during aging and various interventions that promote or maintain protein homeostasis beneficially affect lifespan in model organisms^3-5^. During stressful conditions, the maintenance of protein homeostasis by cellular stress response pathways is an essential feature of cellular integrity and organismal fitness. Internal and external stimuli trigger evolutionarily conserved cellular stress pathways such as the heat shock response (HSR), organelle-specific stress response pathways such as the endoplasmic reticulum or mitochondrial unfolded protein responses (ER-UPR/mito-UPR) and the Integrated Stress Response (ISR). Multiple lines of evidence show that longevity ultimately relies on the fidelity of cellular stress response mechanisms^6^.

The biological function of the ISR is to restore cellular homeostasis upon stress. The activation of the ISR relies on the eukaryotic initiation factor 2 (eIF2) kinases: the heme-regulated inhibitor (HRI), protein kinase R (PKR), general control nonderepressible 2 (GCN2), and PKR-like endoplasmic reticulum kinase (PERK). They are activated, respectively, by iron deficiency, viral infection, amino acid deprivation and accumulation of misfolded protein in the ER. The kinases converge on the phosphorylation of the α subunit of eIF2. eIF2 is a key regulator of translation initiation, the limiting step of protein synthesis^7^. For translation initiation to occur, the eIF2.GTP.tRNA^met^ ternary complex together with other initiation factors and the 40S ribosomal subunit forms the 43S pre-initiation complex. The 43S complex binds to the 5’-cap structure and scans along the mRNA until it recognizes the AUG start codon. Then, GTP hydrolysis releases eIF2 and other initiation factors from the mRNA-40S-complex, allowing the 60S ribosomal sub-unit to bind and proceed to elongation^8^. The exchange of GDP to GTP is necessary for recycling of eIF2 back to its active form and for further rounds of translation initiation. This exchange is catalyzed by the heterodecameric guanine nucleotide exchange factor eIF2B. The phosphorylation of eIF2α at serine 51 by the stress sensitive kinases represents the core event of the ISR.

Phospho-eIF2α is a strong inhibitor of eIF2B leading to attenuated ternary complex formation and therefore to a reduction of 5’-cap-dependent protein synthesis. Decreasing the abundance of the ternary complex paradoxically de-represses translation of specific mRNAs that are regulated by upstream open reading frames (uORFs) such as *ATF4, ATF5*, and *CHOP*. While the ISR and translation initiation are finely balanced to provide robustness during acute challenges to protein homeostasis, the role of this pathway during aging and in longevity remains largely unexplored.

Forward genomic screens in *C. elegans* have shed light on numerous pathways whose activity extend lifespan^9,10^. These approaches used systematic mRNA knockdown, and did not have the resolution to investigate consequence of other genetic alterations, including gain-of-function mutations, in longevity. Unbiased forward screens using chemical mutagenesis coupled with whole genome sequencing are a powerful tool to reveal new longevity loci. We therefore set out to perform a large-scale mutagenesis screen for increased survival in *C. elegans*.

## Results

### The ISR is a regulator of longevity and protein homeostasis in *C. elegans*

To identify novel modulators of the aging process, we optimized an unbiased forward longevity genetic screen^11,12^ (Fig. 1a). The conditionally sterile CF512 strain *fer15(b26) II*; *fem-1(hc17) IV* was mutagenized with 0,3% ethyl methanesulfonate. Of 28000 tested genomes, 318 mutant strains showed increased maximum lifespan and after full demographic analyses (Fig. 1b), we sequenced 101 genomes of mutants with a mean lifespan extension of at least 18%. Single-nucleotide polymorphism mapping revealed potential longevity variants in genes that control eIF2. We found two independent alleles in *ppp-1*/eIF2Bγ, one mutation in *gcn-2*/GCN-2 and one mutation in *pek-1*/PERK (Fig. 1b, Extended Data Fig. 1a). These results suggest a link between ISR regulation (Fig. 1d) and *C. elegans* longevity. Outcrossed *ppp-1*(*wrm10*) and *ppp-1*(*wrm15*) alleles extended *C. elegans* lifespan by 20% (Fig. 1e and Extended Data Table 1). Furthermore, CRISPR/Cas9 generated mutants with identical substitutions confirmed the longevity (Fig. 1f). The outcrossed *gcn-2(wrm4)* and *pek-1(wrm7)* mutants as well as the *gcn-2(wrm4)*;*pek-1(wrm7)* double mutant were long-lived (Fig. 1g).

**Fig. 1.**
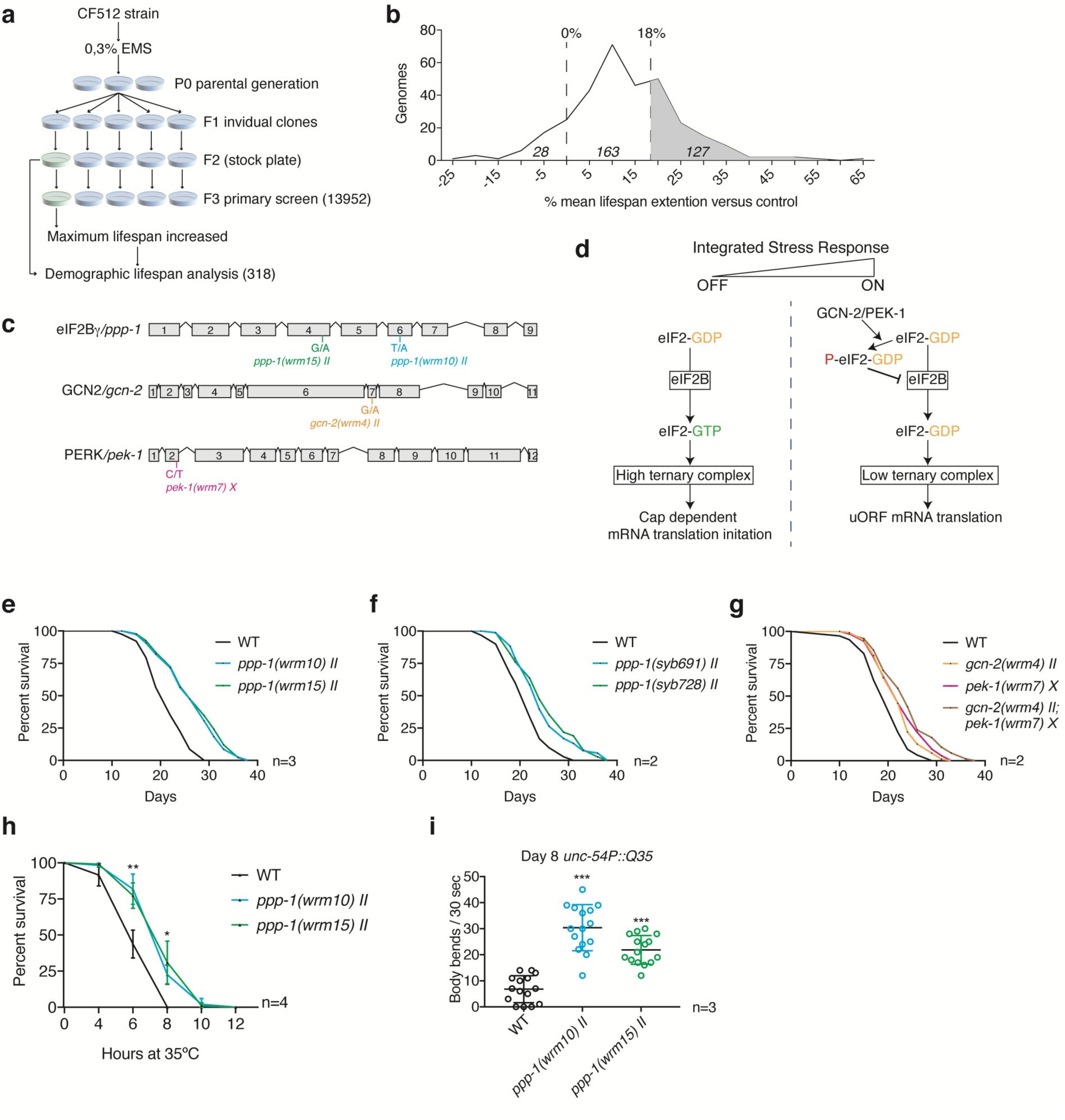
Unbiased forward longevity screen in *C*. *elegans* identifies mutations in ISR components. **a**, Screening strategy. **b**, Percent of mean lifespan extension (compared to temperature sensitive sterile CF512 control) as a function of the number of genomes tested. **c**, Schematic representation of identified ISR genes and corresponding longevity alleles. **d**, Cartoon depiction of the ISR. **e**, Survival of outcrossed *ppp-1*(*wrm10*) and (*wrm15*) mutants compared to WT controls (n=3). **f**, Survival of CRISPR/Cas9-generated *ppp-1* alleles (*syb691*) and (*syb728*) compared to WT controls (n=2). *syb691 corresponds to wrm10 and syb728 to wrm15*. **g**, Survival of outcrossed *gcn-2(wrm4), pek-1(wrm7)* and double *gcn-2(wrm4);pek-1(wrm7)* mutants compared to WT controls (n=2). **h**, Thermotolerance assays of day 1 *ppp-1* mutant worms show significantly increased survival during heat stress compared to WT (error bars represent means ±SD, two-way ANOVA Dunnett’s post hoc test with **p<0.01 and *p<0.05 versus WT controls; n=4). **i**, Motility assays using day 8 WT and *ppp-1* mutants with *unc-54P*-driven muscle-specific expression of polyQ35-YFP fusion protein (error bars represent means ±SD, one-way ANOVA Dunnett’s post hoc test with ***p<0.001 versus WT controls; n=3). See Extended Data Table 1 for lifespan statistics. See Extended Data Table 2 for statistics on thermotolerance assays.

We further characterized *ppp-1* mutants using proteotoxic challenges. Upon heat shock, *ppp-1* mutants showed enhanced survival compared to WT animals (Fig. 1h; Extended Data Table 2). Expression of fluorescently tagged polyglutamine (polyQ35) stretches in the muscle^13^ results in a drastic decrease of motility (Fig. 1i). Strikingly, *ppp-1* mutants were protected from polyQ35 toxicity (Fig. 1i). Together, these results demonstrate that *ppp-1* mutations extend lifespan and protect from proteotoxicity.

### Gcn(-) mutations extend *C. elegans* lifespan

We used RNAi to investigate the effect of *ppp-1* silencing. *ppp-1* knockdown did not affect the lifespan of WT animals (Fig. 2a). This was unexpected as silencing eIF2Bδ reduced protein synthesis and extends *C. elegans* lifespan^14^. Instead, *ppp-1* RNAi abolished longevity and heat resistance of both *ppp-1* mutants (Fig. 2a; Extended Data Fig. 2a) and heterozygous *ppp-1* mutants were long-lived (Fig. 2b). These observations suggest that *ppp-1* mutations are genetically dominant. Activation of *ppp-1*, hence of the eIF2B complex, would reduce the ISR upon stress. To test this hypothesis, we monitored the uORF-regulated translational activation of the worm homolog of GCN4/ATF4, *atf-5*. While DTT treatment significantly increased reporter expression in the WT, both *ppp-1* alleles showed a blunted *atf-5* response during stress (Fig. 2c; Extended Data Fig. 2b). The class of general control non-derepressible (Gcn) mutants in yeast are unable to de-repress the translation of the uORF-regulated transcription factor GCN4/ATF4 upon amino acid starvation^15^. The inability to derepress GCN4/ATF4 mimics a state of inactivated ISR. Taken together, the dominant *ppp-1* mutations show the Gcn(-) phenotype.

**Fig. 2.**
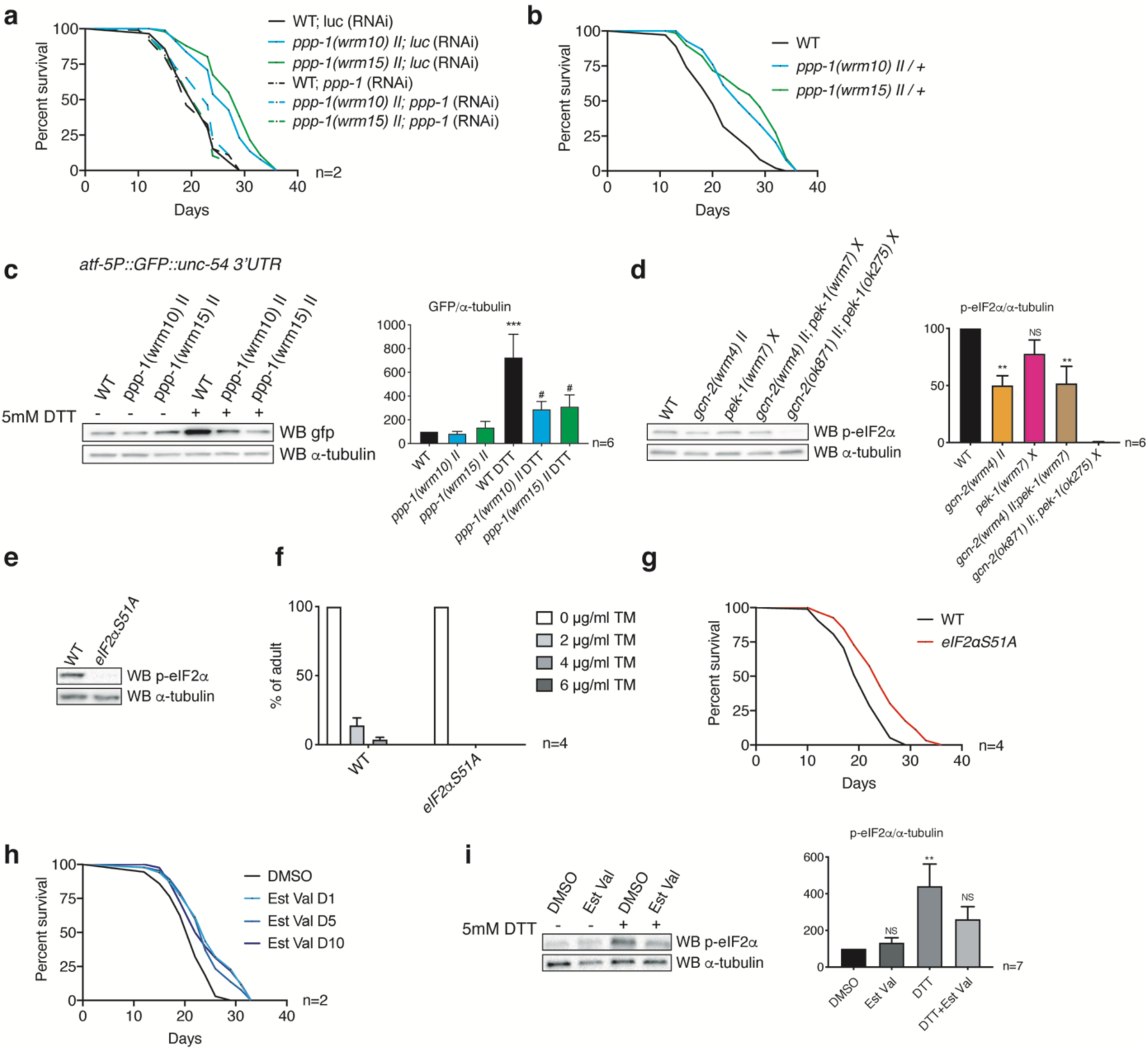
ISR inhibition mediated by Gcn(-) mutations extends *C*. *elegans* lifespan. **a**, Survival of WT and *ppp-1* mutants upon RNAi treatment targeting *ppp-1* and control *luciferase* (*luc*) (n=2). **b**, Survival of heterozygous *ppp-1 mutants* compared to WT control animals. **c**, Representative Western blot of day 1 WT and *ppp-1* mutants in the *atf-5P*::GFP::*unc-54* 3’UTR reporter background treated with 5 mM DTT for 2 hours, using anti-GFP and anti-α-tubulin antibodies. GFP levels were normalized to α-tubulin (error bars represent means +SEM, one-way ANOVA Tukey’s post hoc test, ***p<0.001 versus WT(-DTT), #p<0.05 versus WT(+DTT); n=6). **d**, Representative Western blot of day 1 worms of indicated genotypes detecting phospho-eIF2α (Ser51) and α-tubulin. Levels of phospho-eIF2α were normalized to α-tubulin (error bars represent means +SEM, one-way ANOVA Dunnett’s post hoc test, **p<0.01 versus WT, NS=not significant versus WT; n=6). **e**, Western blot of day 1 *eIF2*α*S51A* mutants using anti-phospho-eIF2α (Ser51) and anti-α-tubulin antibodies. **f**, Developmental tunicamycin (TM) resistance assay of WT and *eIF2*α*S51A* mutants treated with indicated TM concentrations (error bars represent means +SEM, two-way ANOVA Sidak’s post hoc test; n=4). **g**, Survival of *eIF2*α*S51A* mutants compared to WT control animals (n=4). **h**, Survival of WT worms treated with 1% DMSO (control) or 20µM Estradiol Valerate (Est Val) from day 1 (D1), day 5 (D5) or day 10 (D10) (n=2). **i**, Representative western blot of day 1 worms treated with 1% DMSO (control) or 20µM Est Val. Worms were incubated without (-) or with 5mM DTT (+) for 2h. Levels of phospho-eIF2α (Ser51) were normalized to α-tubulin (error bars represent means +SEM, one-way ANOVA Dunnett’s post hoc test, **p<0.01 versus WT(-DTT), NS=not significant versus WT(-DTT); n=6). See Extended Data Table 1 for lifespan statistics.

Next, we tested whether the *gcn-2(wrm4)* and *pek-1(wrm7)* mutants also belong to the Gcn(-) class. The *gcn-2(wrm4)* mutant displayed a 50% reduction of baseline eIF2α phosphorylation suggesting that this mutant can be classified as Gcn(-) (Fig. 2d). The reduction of eIF2α phosphorylation in the *pek-1(wrm7)* mutant did not reach significance. To mechanistically address whether Gcn(-) mutations lead to longevity, we engineered a phospho-defective *eIF2*α*S51A* mutant (*eIF2*α*(syb1385))*, abolishing the ISR (Fig. 2e). Homozygous *eIF2*α*S51A* mutants were viable and displayed regular pharyngeal pumping rates (Extended Data Fig. 2c), generation time (Extended Data Fig. 2d), and brood size (Extended Data Fig. 2e). Importantly, *eIF2*α*S51A* mutants were hypersensitive to ER stress induced by tunicamycin, likely because phosphorylation of eIF2α by the *pek-1*/PERK kinase is required to promote the ER stress response and survival (Fig. 2f). Notably, *eIF2*α*S51A* mutants showed a robust lifespan extension compared to WT animals demonstrating that Gcn(-) mutations lead to longevity in *C. elegans* and that the genetic inhibition of the ISR can extend lifespan (Fig. 2g). Consistently, *eIF2*α*S51A* mutants were heat resistant (Extended Data Fig. 2f). Finally, we assessed survival during pharmacological ISR inhibition. For this, we used a set of compounds that were previously described as UPR modulators in worms^16^. Estradiol valerate reduced GFP induction of the *atf-5* reporter during tunicamycin treatment whereas propafenone hydrochloride further elevated GFP expression (Extended Data Fig. 2g). Consistent with the Gcn(-) phenotype, estradiol valerate significantly extended *C. elegans* lifespan (Fig. 2h) and suppressed eIF2α phosphorylation upon DTT treatment (Fig. 2i). Surprisingly, treatments initiated at day 5 or day 10 of adulthood equally increased survival (Fig. 2h) suggesting that late ISR inhibition might be sufficient to promote lifespan extension. ISR induction with propafenone hydrochloride shortened lifespan (Extended Data Fig. 2h).

### Gcn(-) longevity is independent of attenuated translation

As eIF2B is a key regulator of translation initiation, we monitored protein synthesis in *ppp-1* mutants. First, we measured the levels of incorporated radioactive methionine of day 1 adult animals and did not observe any differences between WT animals and *ppp-1* mutants (Fig. 3a). To corroborate these results, we used surface sensing of translation (SUnSET) as an alternative measurement of protein synthesis rates. This technique is based on the incorporation of puromycin into newly synthesized peptides followed by the detection of the labelled peptides with monoclonal antibody^17^. No changes in protein synthesis were observed between *ppp-1* mutants and WT animals whereas control *rsks-1/*S6K mutants showed a drastic reduction of puromycin-labelled peptides (Fig. 3b). Finally, we performed polysome profiling to evaluate the distribution of the ribosomal subunits and complexes after separation on sucrose gradients. We found no differences in the overall ribosome distribution and abundance (Fig. 3c-d). Likewise, we found no differences in polysome abundance at day 1 of adulthood between WT animals and the *eIF2*α*S51A* mutant (Fig. 3e-f). Since eIF2 activity is regulated by phosphorylation, we also evaluated the level of phospho-eIF2α on day 1 and day 6 of adulthood. We found that the phosphorylation of eIF2α was increased upon aging in WT animals (Extended Data Fig. 3a). However, we did not observe any differences between *ppp-1* mutants or WT control at day 1 or day 6. Together, our results support the idea that the longevity of *ppp-1* mutants is uncoupled from reduced protein synthesis.

**Fig. 3.**
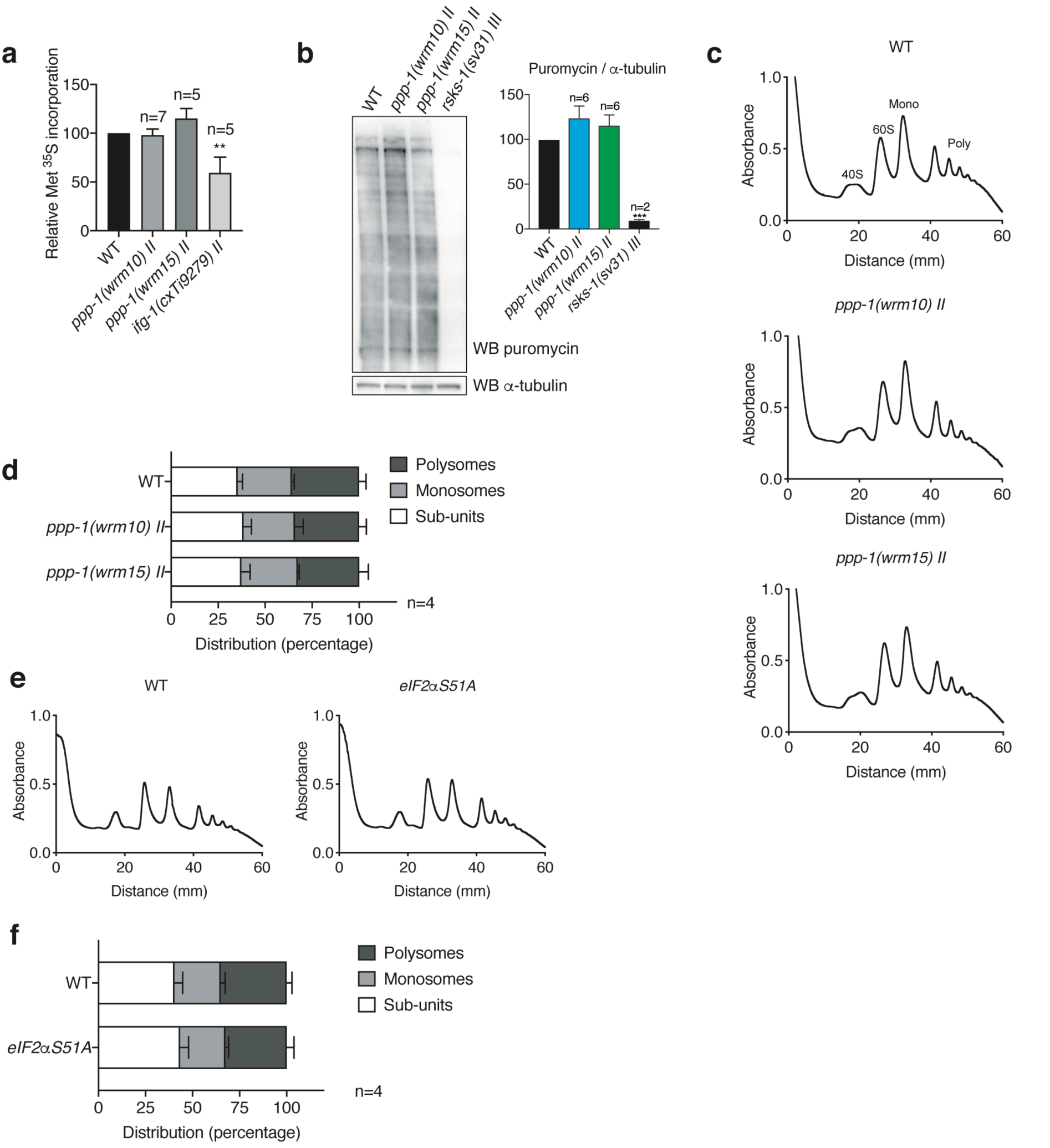
Gcn(-) mutants show no changes in overall protein biosynthesis. **a**, ^35^S-methionine labelling of day 1 WT worms, *ppp-1* mutants and control *ifg-1(cxTi9272)* mutants (error bars represent means +SEM, one-way ANOVA Dunnett’s post hoc test with **p<0.01 versus WT; biological replicates (n) as indicated in the figure). **b**, Puromycin incorporation followed by Western blot analysis using antibodies detecting puromycin and α-tubulin in day 1 WT animals, *ppp-1* mutants, and control *rsks-1(sv31)* mutants (error bars represent means +SEM, one-way ANOVA Dunnett’s post hoc test with ***p<0.001 versus WT; biological replicates (n) as indicated in the figure). **c, d**, Polysome profiling and quantification of day 1 WT and *ppp-1* animals. Quantification represents the relative abundance of ribosomal subunits (40S, 60S), monosomes (mono) and polysomes (poly) (error bars represent means +SD, two-way ANOVA Dunnett’s post hoc test; n=4). **e, f**, Polysome profiling and quantification of day 1 WT worms and *eIF2αS51A* mutants (error bars represent means +SD, two-way ANOVA Dunnett’s post hoc test; n=4).

### *kin-35* translation is required for Gcn(-) longevity

As we did not observe any changes in global protein synthesis, we asked whether translational efficiency of specific mRNAs could be causative for the lifespan extension of the *ppp-1* animals. We compared the ratio of polysome-associated mRNAs (>3 ribosomes/mRNA) normalized to total mRNA levels between WT and *ppp-1* animals (Fig. 4a). We found a significant de-enrichment of 336 mRNAs and an enrichment for 72 mRNAs in *ppp-1* polysome fractions (Fig. 4b). GO Term analysis revealed an enrichment for genes involved in phosphorylation (Fig. 4c).

**Fig. 4.**
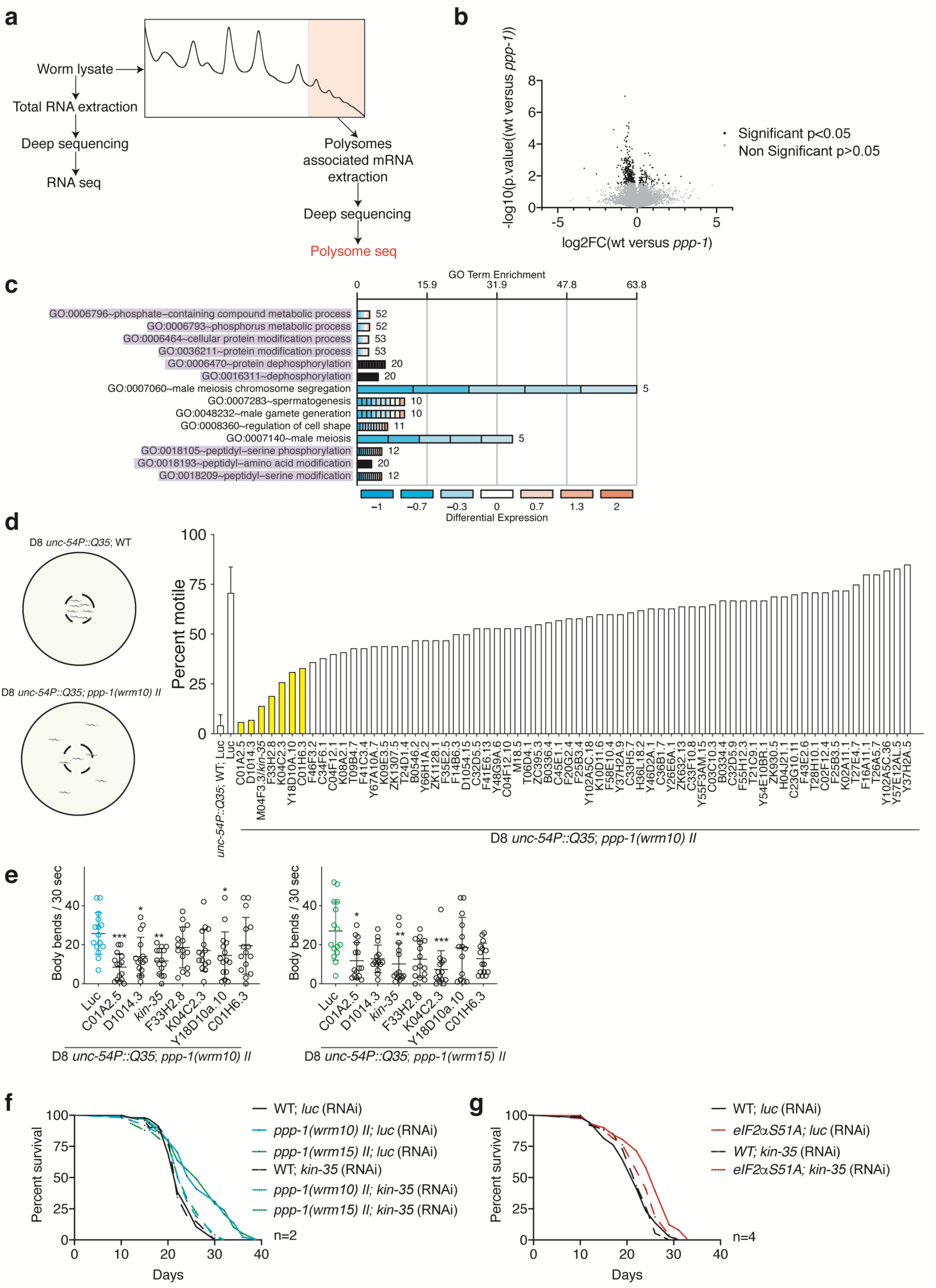
*kin-35* translation is required for longevity of Gcn(-) mutants. **a**, Polysome sequencing strategy. **b**, Volcano plot of polysome-associated mRNAs normalized to total mRNA levels between WT and *ppp-1* mutants. All displayed mRNAs were found in both *ppp-1* mutants. Mean p-values and mean log-2 fold change of both *ppp-1* mutants were used (Student’s t-test, significance is reached for p<0.05). FC = fold change. The full dataset can be found in Extended Data Table 3. **c**, DAVID gene ontology (GO) analysis of significantly changed mRNAs shown in (B). Processes involved in phosphorylation are highlighted in purple. **d**, Selective RNAi screen for suppressors of *ppp-1(wrm10)* polyQ35 motility. For more reliability, assays of WT polyQ35 and *ppp-1(wrm10)* polyQ35 on *luc* RNAi were performed four times (error bars represent means +SD). **e**, Motility assays of day 8 WT polyQ35 and *ppp-1* polyQ35 mutants after indicated RNAi treatments (error bars represent means ±SD, one-way ANOVA Dunnett’s post hoc test with *p<0.05, **p<0.01 and ***p<0.001 versus *luc* control). **f**, Survival of WT and *ppp-1* mutants upon *kin-35* and control *luc* RNAi knockdown (n=2). **g**, Survival of WT and *eIF2αS51A* mutants upon RNAi knockdown of *kin-35* and control *luc* (n=4). See Extended Data Table 1 for lifespan statistics.

Several studies have demonstrated that translation efficiency of specific mRNAs is a key regulator of lifespan under different longevity paradigms in *C. elegans*^*18,19*^. Therefore, we hypothesized that some of the enriched mRNAs define *ppp-1* phenotypes. We used resistance to polyQ35 proteotoxicity of *ppp-1* animals as a proxy for longevity and knocked down the candidate mRNAs in *ppp-1(wrm10)* mutants with RNAi. At day 8 of adulthood, all polyQ35 transgenic animals were paralyzed. polyQ35;*ppp-1(wrm10)* animals remained motile and were screened for suppressors (Fig. 4d). We found seven RNAi clones that abolished the motility of the *ppp-1* mutants by at least 50%. Motility assays quantifying body bending in liquid validated these results in both *ppp-1* mutant alleles (Fig. 4e). Knockdown of candidate genes C01A2.5 and M04F3.3 showed no motility reduction in WT animals (Extended Data Fig. 4a) but significantly decreased motility in both *ppp-1* mutants. Lifespan analyses next showed full suppression of *ppp-1* and *eIF2*α*-S51A* longevity (Fig. 4f-g) upon M04F3.3 knockdown. M04F3.3 encodes a predicted kinase with yet unknown function in the worm that we termed *kin-35*. qPCR analysis confirmed that *kin-35* mRNA association with polysomes was enhanced in *ppp-1* mutants without increased allover abundance (Extended Data Fig. 4b). Together, these data suggest that increased translation of *kin-35* mRNA is required for *ppp-1* longevity. C01A2.5 also significantly reduced *ppp-1* longevity but also shortened WT lifespan suggesting general toxicity (Extended Data Fig. 4c). Our results demonstrate that selective translation of *kin-35* is required for lifespan extension and increased protein homeostasis in Gcn(-) mutants.

## Discussion

Through an unbiased genomic screen for longevity in *C. elegans*, we identified the ISR as a longevity pathway. We provide evidence that genetic inhibition of the ISR via Gcn(-) class mutations or via pharmacological treatment extend lifespan. Gcn(-) mutations attenuate the stress-induced expression of uORF-regulated genes such as ATF4/GCN4, inhibiting the ISR^15^. Mutations that reduce or abolish eIF2α phosphorylation, as in the partial *gcn-2* loss-of-function and the eIF2α-S51A mutants analysed in this study therefore belong to the Gcn(-) class. We also classified the dominant eIF2Bγ/*ppp-1* alleles as Gcn(-) mutations as they reduced uORF regulated *atf-5* expression under stress. eIF2B subunits have been identified carrying Gcn(-) mutations in yeast^20^. Upon phosphorylation, eIF2α inhibits eIF2B^21^ and mutations in eIF2Bβ/GCD7 and eIF2Bγ/GCD2 render eIF2B insensitive to its inactivation by phosphorylated eIF2α^20,22^. These eIF2B variants are protected from inhibition during the ISR. The eIF2Bγ/*ppp-1* mutants we found might have similar features regarding regulation by phosphorylated eIF2α and thus showed decreased ISR activity. In line with the lifespan extension, eIF2B and *eIF2*α*S51A* mutants displayed improved protein homeostasis, essential for cellular and organismal health. Reduced mRNA translation is associated with longevity^23-27^. The long-lived eIF2B and *eIF2*α*S51A* mutants showed maintained translation rates. Hence, longevity did not involve reduced protein biosynthesis. Instead, translational efficiency of specific mRNAs was altered; selective translation of the kinase KIN-35 was required for the longevity of Gcn(-) mutants.

### What is the link between Gcn(-) mutations and lifespan extension?

The regulation of translation initiation and the ISR are intimately linked. Our data suggest that a shift in the translatome, and not the loss of the ISR *per se*, is responsible for extending lifespan. Long-lived *daf-2*/insulin receptor mutants show changes in their translatome^28^ and the extended lifespan of *daf-2;rsks-1*/S6K double mutants is mediated by the selective translational repression of the cytochrome *cyc-2*.*1*^18^. Our study shows that Gcn(-) mutations change translational efficiency of specific mRNAs that are required for the observed lifespan extension. This is in line with a regulation of aging at the level of mRNA translation. While it is not understood how *kin-35* mRNA is selectively recruited to polysomes, our data suggest that upregulation of KIN-35 constitutes a switch that enhances robustness through phosphorylation. This is supported by the analysis of polysome associated mRNAs that points to a broader change in the cellular dynamics of phosphorylation and dephosphorylation.

A number of interventions that extend mouse lifespan show elevated ATF4 expression^29^ and ATF4 is linked to lifespan extension via FGF21 in mice^30^. Additionally, GCN4 is required in yeast to extend lifespan when translation is inhibited suggesting a beneficial effect of activated ISR for longevity^31^. Further, pharmacological ISR activation is protective in a Huntingtin mouse model^32^. Nevertheless, deregulated activation of the ISR has also been correlated with cancer and diabetes^33,34^. The ISR is activated in neurogenerative disorders, traumatic brain injury, and Down syndrome^35-38^. Although the role of the ISR in longevity is thus unclear and is very likely to differ between cell types, no studies have yet formally tested how direct modulation of the ISR affects mammalian survival. Our data show that reducing or fully abrogating the ISR in Gcn(-) mutants extended *C. elegans* lifespan. While the ISR is clearly required to cope with acute stress, the translatome changes in Gcn(-) mutants appear to support robustness and protein homeostasis.

Pathological conditions associated with an increased ISR can be treated by reducing eIF2α phosphorylation or by interfering with the inhibition of eIF2B. Deletion of eIF2α kinases prevents pathology in a mouse model for Alzheimer’s disease^35^ and PKR knockout enhances cognitive function in a mouse model for Down syndrome^37^. This suggests a causal role of the ISR in these age-associated diseases. Further, memory is enhanced in mice heterozygous for the eIF2αSer51Ala mutation^39^. Pharmacological inhibition of the ISR is possible using the small molecule ISRIB, which enhances memory, prevents neurodegeneration in prion disease, and reverses memory defects associated with traumatic brain injury^36,38,40^. Mechanistically, ISRIB stabilizes and activates eIF2B, which counters the effects of eIF2α phosphorylation^41,42^. In all, these data converge with the enhanced survival and robustness we observed in the Gcn(-) *C. elegans* mutants. Pharmacological inhibition of the ISR might be a promising therapeutic approach to modulate the ageing process.

## Methods

### *C. elegans* strains and culture

All *C. elegans* strains were maintained at 20°C on nematode growth medium (NGM) agar plates seeded with the *Escherichia coli* (*E. coli*) strain OP50, unless indicated otherwise^43^. To provide an isogenic background in all mutant strains, they were outcrossed against the wildtype Bristol N2 strain. All strains used in this study are listed in Extended Data Table 4, including outcrossing information and source. Genotyping primers used in this study are listed in Extended Data Table 5. The strains *ppp-1(syb728) II* and *ppp-1(syb691) II* were generated by SunyBiotech (China) using CRISPR/Cas9; the correct sequence was verified by PCR and Sanger sequencing (Eurofins Genomics, Germany).

### Unbiased forward longevity screen

The longevity screen was performed with the temperature sensitive sterile strain CF512 *fer-15(b26) II; fem-1(hc17) IV*. L4 larvae were exposed to 0,3% ethyl methane sulfonate (EMS, Sigma) in M9 buffer for 4 h at room temperature. After recovery overnight, young P0 adult animals were transferred to new plates. Singled F1 progeny were allowed to lay eggs overnight. In the next generation, singled F2 progeny were allowed to lay eggs for 16 h. After egg-laying, F2 worms were stocked at 15°C. F3 eggs were heat shocked at 25°C for 48 h to induce sterility and adult animals were scored twice a week for preliminary lifespan analysis. Mutants that outlived the non-mutagenized control by 20% (maximum lifespan) were selected for regular demographic lifespan analyses to confirm the longevity phenotype. After the lifespan assays, mutants with a mean lifespan extension above 18% compared to non-mutagenized CF512 controls were selected for whole genome sequencing.

### Mutant mapping and sequence analysis

Genomic DNA of select long-lived strains was prepared using the QIAGEN Gentra Puregene Kit. Whole genome sequencing was conducted on the Illumina HiSeq2000 platform. Paired-end 100 bp reads were used; the average coverage was larger than 16-fold. Sequencing outputs were analyzed using the CloudMap Unmapped Mutant Workflow pipeline on Galaxy^44^. The WS220/ce10 *C. elegans* assembly was used as reference genome.

### Lifespan assays

Gravid day 1 adults were allowed to lay eggs for 5 h. The offspring was used for lifespan analysis. The L4 stage was defined as day 0 and more than 100 worms were used per strain and condition. Worms were kept at 20°C on NGM plates seeded with OP50 *E. coli* at all times. The animals were transferred every second day to fresh plates until they reached the post-reproductive stage. Scoring was performed every second day by monitoring (touch-provoked) movement and pharyngeal pumping. Animals in all RNAi lifespan assays were treated with RNAi from the young adult stage on and kept on NGM plates seeded with HT115 *E. coli* bacteria expressing control *luciferase* or candidate RNAi clones throughout the experiment. Animals in all lifespan assays on estradiol valerate (Est Val, Sigma) or propafenone hydrochloride (Propa, Sigma) were transferred at the L4 stage to NGM plates containing 1% DMSO (Sigma) and 20 µM Est Val/Propa or to control plates with 1% DMSO only. Lifespan assays of heterozygous animals were performed on F1 hermaphrodites after crossing of mutant hermaphrodites to wildtype male animals. In all lifespan experiments, worms that had undergone internal hatching, vulval bursting, or worms crawling off the plates were censored. Throughout the experiment, strain and/or treatment was unknown to researchers. Data were assembled on completion of the experiment. Statistical analyses were performed with the Mantel-Cox log rank method in Prism (Version 8.2.0).

### Thermotolerance assays

After an egg-lay, synchronized day 1 animals were transferred to 6 cm NGM plates containing OP50 and placed at 35°C. Survival was scored for (touch-provoked) movement and pharyngeal pumping every two hours until no survivors were left. Worms with internal hatching, vulval bursting, and worms crawling off the plates were censored. Throughout the experiment, strain and/or treatment was unknown to the researcher. Unless stated otherwise, at least 3 independent experiments were performed, error bars represent means ±SD and assays were analyzed by two-way ANOVA, Dunnett’s or Sidak’s post hoc test as indicated.

### Motility assays in *unc-54P::Q35:YFP* background

Animals carrying the *unc-54P::Q35:YFP* (polyQ35) transgene were grown on NGM plates seeded with OP50. For RNAi experiments, they were transferred at the L4 stage to plates seeded with HT115 bacteria expressing *luciferase* or candidate RNAi clones. On day 8 of adulthood, motility was tested by transferring single worms to M9, where they were allowed to acclimatize for 30 sec, followed by the counting of body bends over 30 sec. At least 12 worms were scored per experiment, genotype and/or treatment. Throughout the experiment, strain and/or treatment was unknown to the researcher. Unless stated otherwise, at least 3 independent experiments were performed, error bars represent means ±SD and assays were analyzed by one-way ANOVA, Dunnett’s post hoc test.

### RNAi experiments

For RNAi mediated knockdown of specific genes, HT115 bacteria carrying vectors for dsRNA of the target gene under a promotor inducible by isopropyl β-D-1-thiogalactopyranoside (IPTG) and an ampicillin resistance were used. Bacteria were seeded on NGM plates containing 100 µg/µL ampicillin (Merck Millipore) and 1 mM IPTG (Roth). After egg-lay, worms were grown on regular NGM plates seeded with OP50 bacteria until the L4 stage and then transferred to RNAi plates. RNAi against *luciferase* was used as nontargeting control. All RNAi clones were obtained from the Ahringer and Vidal RNAi libraries^45,46^. Clones were validated by plasmid purification (QIAprep Spin Miniprep Kit, Qiagen) and sequencing using the L4440 seq RV primer.

### Induction of endoplasmic reticulum stress with dithiothreitol

Endoplasmic reticulum (ER) stress was induced by incubation of worms in dithiothreitol (DTT, Sigma). For the DTT treatment, an overnight culture of OP50 bacteria was 10-fold concentrated in S-basal medium. Worms were transferred into 250 µL S-basal medium, 200 µL 10-fold concentrated OP50 and 5 µL 1 M DTT diluted in S-basal. The volume was filled up to a total of 1 mL with S-basal (final DTT concentration: 5 mM). Worms were incubated for 2 h at 200 rpm.

### Western blotting

For Western blotting, day 1 worms were collected in M9 and snap frozen in liquid nitrogen. For protein extraction, worms were lysed in Ripa buffer (150 mM NaCl, 1% NP40, 0.5% sodium deoxycholate, 0.1% SDS, 50 mM Tris-HCl, pH 8.0, completed with protease inhibitors), sonicated and spun down. The supernatant was taken to protein quantification by bicinchoninic acid assay (Pierce BCA Protein Assay Kit, Thermo Fisher). Equal amounts of protein were taken to NuPAGE LDS Sample Buffer (4X, ThermoFisher) containing 50 mM Dithiothreitol (DTT). Proteins were then separated by reducing SDS-PAGE and transferred to nitrocellulose membranes (AmershamTM Hybond ECL), followed by blocking with milk or BSA and antibody labelling with specific antibodies to phospho-eIF2α (Ser51) (Cell Signaling), puromycin (Merck Millipore), Living Colors GFP (Clontech) and α-tubulin (Sigma). Immunolabeling was visualized using chemiluminescence kits (ECL, Amersham Bioscience) on a Chemidoc MP Imaging System (Biorad) and analyzed with the ImageLab Software (version 5.2, Biorad). Labeling was quantified with Image J (version 1.51) and Prism (version 8.2.0). For Western blot analyses of compound-feeding experiments, worms were fed after hatching with 20 µM Est Val and 1% DMSO, or 1% DMSO only. ER stress by DTT was induced as described above. For Western blot analysis at day 6 of adulthood (and corresponding day 1 control experiments), worms were transferred to NGM plates containing 10 µM 5-Fluoro-2′-deoxyuridine (Sigma) at the L4 stadium. The collection of the Western blot samples was conducted at the same time for day 1 and day 6 animals. Unless stated otherwise, at least 4 independent experiments were performed, error bars represent means ± SEM and assays were analyzed by one-way ANOVA, Tukey’s or Dunnett’s post hoc test as indicated.

### Developmental tunicamycin resistance assays

For developmental tunicamycin (TM) resistance assays, NGM plates supplemented with 10 µg/mL TM and control plates without TM were used (seeded with OP50 bacteria). 50 to 80 synchronized eggs per genotype and/or condition were added to the plates. Development to the adult stage was scored after 4 or 5 days. Unless stated otherwise, at least 4 independent experiments were performed, error bars represent means ±SEM and assays were analyzed by two-way ANOVA, Sidak’s post hoc test.

### ^35^S-methionine labelling

To monitor translation rates, ^35^S-methionine labelling was performed based on Hansen *et al*., 2007^25^. OP50 bacteria were cultured overnight in LB medium (1 mL/sample) containing 15 μCi of ^35^S-methionine and concentrated 10-fold. Synchronized day 1 worms were added to the mix and incubated for 3 h at room temperature. Worms were washed twice with S-basal and incubated in non-radioactive OP50 (10-fold concentrated). Worms were washed twice with S-basal medium and flash frozen three times in liquid nitrogen. Worm pellets were boiled in 100 μL 1% SDS and centrifuged 2 min at 2000 g to remove cuticles. Supernatants were submitted to trichloroacetic acid precipitation. Protein pellets were neutralized with 20 μL of 0,2 M NaOH. Proteins were solubilized with 180 μL of 8 M urea; 4% chaps; 1% DTT. Protein concentrations were measured using Bradford reagent and ^35^S radioactivity was measured by liquid scintillation. Unless stated otherwise, at least 5 independent experiments were performed, error bars represent means ±SEM and assays were analyzed by one-way ANOVA, Dunnett’s post hoc test.

### Surface sensing of translation (SUnSET), puromycin incorporation

To monitor protein synthesis in a non-radioactive manner using puromycin incorporation and puromycin detection based on Schmidt *et al*., 2009 ^17^, day 1 worms were collected in M9 and once washed into S-basal medium. For the puromycin treatment, an overnight culture of OP50 bacteria was 10-fold concentrated in S-basal medium. Worms were then transferred into 250 µL S-basal medium, 200 µL 10-fold concentrated OP50 and 50 µL 10 mg/mL puromycin diluted in S-basal. The volume was filled up to a total of 1 mL with S-basal (final puromycin concentration: 0,5 mg/mL). Worms were incubated for 3 h at 200 rpm. Afterwards, they were washed 3 times in S-basal and snap-frozen in liquid nitrogen. Worms were kept on ice after the puromycin treatment. Protein extraction and Western blot using anti-puromycin antibody (Merck Millipore) was performed as described before.

### Polysome profiling

For the analysis of translation via polysome profiling based on Ding and Großhans, 2009^47^, synchronized gravid day 1 adults were grown on NGM plated seeded with OP50. Per genotype and replicate, ∼12000 worms were harvested and washed twice with M9, once with M9 supplemented with 1 mM cycloheximide (Sigma) and once with lysis buffer (20 mM Tris pH 8.5, 140 mM KCl, 1.5 mM MgCl2, 0.5% Nonidet P40, 1 mM DTT, 1 mM cycloheximide). Worms were pelleted and resuspended in 350 µL cold lysis buffer supplemented with 1% sodiumdeoxycholate (DOC, Sigma). Resuspended worms were lysed using a chilled Dounce homogenizer. Ribonuclease inhibitor RNasin (Promega) was added to samples used for RNA sequencing or quantitative PCR (qPCR) at a concentration of 0,4 U/µL. Samples were then mixed and incubated on ice for 30 min, followed by a centrifugation step (12000 g, 10 min, 4°C) for clearance. The pellet was discarded and the RNA concentration of the supernatant was estimated by absorbance measurement at 260 nm.

To prepare sucrose gradients, 15% (w/v) and 60% (w/v) sucrose solutions were prepared in basic lysis buffer (20 mM Tris pH 8.5, 140 mM KCl, 1.5 mM MgCl2, 1 mM DTT, 1 mM cycloheximide). Linear sucrose gradients were produced using a Gradient Master (Biocomp). Equivalent amounts of sample (around 400 µg RNA) were loaded on the gradient and centrifuged at 39000 g for 3 h at 4°C, using an Optima L-100 XP Ultracentrifuge (Beckman Coulter) and the SW41Ti rotor. To analyze the sample on the gradient during fractionation, absorbance at 254 nm was measured and recorded (Econo UV monitor EM-1, Biorad) using the Gradient Profiler software (version 2.07). Gradient fractionation was performed from the top down using a Piston Gradient Fractionator (Biocomp) and a fraction collector (Model 2110, Biorad). Gradients were fractionated in 20 fractions of equal volume. In an initial experiment, the ribosomal fractions were validated by analyzing RNA from each fraction via agarose gel electrophoresis. The 18S and 28S rRNA signals were used as indicators for the 40S ribosomal subunit, the 60S ribosomal subunit and fully assembled ribosomes. Quantification of the ribosomal complexes was performed using Image J and statistically analyzed with Prism. Unless stated otherwise, at least four independent experiments were performed, error bars represent means ±SD and assays were analyzed by two-way ANOVA, Dunnett’s post hoc test.

For more precise analysis of ribosomal fractions, they were collected by hand according to their absorbance profile; for RNAseq and qPCR analyses, one fraction for 80S ribosomes and one for polysomes (excluding disomes) was collected per sample. RNA extraction from total lysates and from each fraction was performed using the Direct-zol RNA MicroPrep Kit (Zymo Research) according to the manufacturer’s recommendations.

### Polysome sequencing

For polysome sequencing, monosome extracts, polysome extracts (without disomes), and corresponding total RNA were collected as detailed above. cDNA libraries were generated with ribosomal RNA depletion at the Cologne Center for Genomics and sequenced on the Illumina HiSeq2000 platform.

For data analysis, raw reads from all RNAseq and polysome sequencing replicates were mapped to the *C. elegans* reference genome (ENSEMBL 91) using HISAT2 (v2.1.0)^48^. After guided transcriptome assembly with StringTie (v1.3.4d), transcriptomes were merged with Cuffmerge and quantification was performed with Cuffquant^49^. The analysis for differential gene expression for total, monosomal and polysomal RNA was performed with Cuffdiff (Cufflinks v2.2.1)^50,51^. To analyze the translatome, the abundance of each mRNA in the polysomal fraction was normalized to its abundance in the total input mRNA. Respective normalized values were used to identify changes between different conditions using Student’s t-test. For further analyses, we only included the mRNAs that were found significantly changed in both *ppp-1* mutants. For each mRNA, the mean p-values and the mean log-2 fold change of both *ppp-1* mutants were used. David analysis was performed to identify significantly enriched gene ontology terms^52^.

### Selective RNAi screen for suppressors of *ppp-1* motility

Synchronized worms of the *ppp-1(wrm10)* strain crossed to *mLs133[unc-54P::Q35:YFP]* animals (*ppp-1* polyQ35) and control *mLs133[unc-54P::Q35:YFP]* worms (WT polyQ35) were grown to the L4 larval stadium. Animals were then placed on NGM plates containing 10 µM 5-Fluoro-2′-deoxyuridine (FUDR, Sigma) to inhibit the development of progeny. Plates were seeded with HT115 bacteria expressing selected RNAi clones to knock down specific genes in the nematodes. At day 8 of adulthood, the motility of *ppp-1* polyQ35 as well as WT polyQ35 worms was assessed on *luciferase* control RNAi and 66 RNAi treatments targeting mRNAs enriched in *ppp-1* polysomes. To test motility, 15 worms were picked into the center of a 10 mm circle on an unseeded NGM plate and their ability to leave the circle after one minute was scored. For more reliability, 4 experiments were performed for the control conditions (WT polyQ35 and *ppp-1* polyQ35 on *luciferase* RNAi; error bars represent means ±SD).

RNAi treatments rescuing the *ppp-1* polyQ35 motility phenotype to at least 50% compared to the *ppp-1* polyQ35 control on *luciferase* RNAi were validated by full motility assays (without usage of FUDR) counting body bends over 30 seconds in liquid. In a counter screen, the effect of the RNAi treatments on WT polyQ35 animals was tested. To this end, young worms were treated as described before and the motility on day 6 of adulthood was scored as described above. If motility of WT polyQ35 worms treated with RNAi against candidate mRNAs was significantly lower compared to animals treated with *luciferase* RNAi, candidates were excluded from further analysis.

### Worm imaging

For worm imaging, animals were arranged in stacks on unseeded NGM plates and kept on ice. Images were taken with a fluorescence microscope (Leica M165FC) and a camera (Leica DFC 3000G). Images were taken and analyzed with the Leica Application Suite X (Version 3.4.1.17822), scale bar as indicated in the figures.

### Compound screen

To identify compounds inhibiting the ISR, synchronized *atf-5P*::GFP::*unc-54* 3’UTR) L4 animals were transferred to NGM plates without or with 4 µg/mL tunicamycin (TM). Furthermore, plates were supplemented with 1% DMSO (Sigma) as control, or with 1% DMSO and 20 µM estradiol valerate (Sigma), ISRIB (Sigma), GSK2606414 (Calbiochem), propafenone hydrochloride (Sigma), azadirachtin (Sigma) or estriol (Sigma), respectively. Day 1 animals were analyzed by fluorescent microscopy as described above.

### Pharyngeal pumping

Pharyngeal pumping rates of synchronized animals were measured at day 1 of adulthood by counting pharyngeal contractions per worm during 30 sec. Per experiment and genotype, at least 15 worms were analyzed. Throughout the experiment, strain and/or treatment was unknown to the researcher. Error bars represent means ±SD.

### Generation time

For generation time assays, synchronized eggs were allowed to develop to adult worms on single plates until they laid the first egg, which was defined as generation time. After 55 h, animals were scored every hour with 15 worms being analyzed per experiment and genotype. Throughout the experiment, strain and/or treatment was unknown to the researcher. Error bars represent means ±SD.

### Brood size assays

For brood size assays, synchronized L4 worms were placed on individual NGM plates seeded with OP50 bacteria. Worms were transferred to fresh plates every 24 h until no more eggs were laid. The number of viable progenies on each plate was counted and summed up per individual parental worm. Per experiment, genotype and/or condition, at least 15 parental worms were analyzed. Error bars represent means ±SD.

### qRT-PCR (qPCR)

For qPCR analyses, day 1 worm samples or indicated samples from ribosome profiling were collected in TRI Reagent (Zymo) and frozen in liquid nitrogen. RNA extraction was performed using the Direct-zol RNA MicroPrep Kit (Zymo Research) according to the manufacturer’s recommendations, followed by cDNA synthesis (iScript cDNA Synthesis Kit, BioRad). qPCRs were performed using Power SYBR Green PCR Master Mix (Applied Biosystems) on a ViiA 7 Real-Time PCR System (Applied Biosystems). Primers for the gene M04F3.3 / *kin-35* were used (Forward CGGTTGAATATTGGTGAGGAGGTT; reverse GCCACCATGATCTCTCTTTCAATCT). Primers for the gene *act-1* were used as internal control (Forward CTACGAACTTCCTGACGGACAAG; reverse CCGGCGGACTCCATACC). Unless stated otherwise, at least three independent experiments were performed, error bars represent means ±SEM and assays were analyzed by two-way ANOVA, Tukey’s post hoc test.

### Statistical analysis

Unless stated otherwise, results are presented as means +/±SD or means +/±SEM. Unless noted otherwise, statistical tests were performed using one-way or two-way ANOVA with Sidak’s, Dunnet’s or Tukey’s multiple comparison test. Significance levels are *p<0.05, **p<0.01, and ***p<0.001 versus WT control unless otherwise noted. Experiments were carried out with at least three biological replicates unless noted otherwise.

## Acknowledgements

We thank all Denzel laboratory members for helpful discussions. We thank the Caenorhabditis Genetics Center (CGC) and Dr. T. Keith Blackwell for worm strains. We thank F. Metge, S. Templer and J. Boucas and all members of the bioinformatics core facility at MPI AGE. We thank the Cologne Center for Genomics for sequencing.

## Author contributions

M.D., L.E.W., and M.S.D. conceived the study. All experiments were performed by M.D., L.E.W., and R.B. The manuscript was written and edited by M.D., L.E.W., and M.S.D.

## Data availability

The RNA sequencing data in this publication have been deposited in NCBI’s Gene Expression Omnibus and are accessible through GEO Series accession number GSE144607 (https://www.ncbi.nlm.nih.gov/geo/query/acc.cgi?acc=GSE144607). All other data is available in the main text or the supplementary materials.

