## Supplementary Information for "eIF2B extends lifespan through inhibition of the integrated stress response"

This file includes:

Extended Data Figures 1 to 4
Extended Data Tables 1 to 5

**a**

| Chromosome | Position | Nucleotide substitution | Gene name | AA substitution | AA position | Allele name |
|---|---|---|---|---|---|---|
| II | 6966468 | T/A | *ppp-1* | N/I | 295 | *ppp-1(wrm10) II* |
| II | 6966802 | G/A | *ppp-1* | L/F | 216 | *ppp-1(wrm15) II* |
| II | 13088136 | G/A | *gcn-2* | D/N | 1217 | *gcn-2(wrm4) II* |
| X | 11414450 | C/T | *pek-1* | D/N | 81 | *pek-1(wrm7) X* |

**Extended Data Fig. 1 | *C. elegans* mutations clustering in the ISR identified through an unbiased forward longevity screen. a**, Detailed overview of identified longevity alleles.

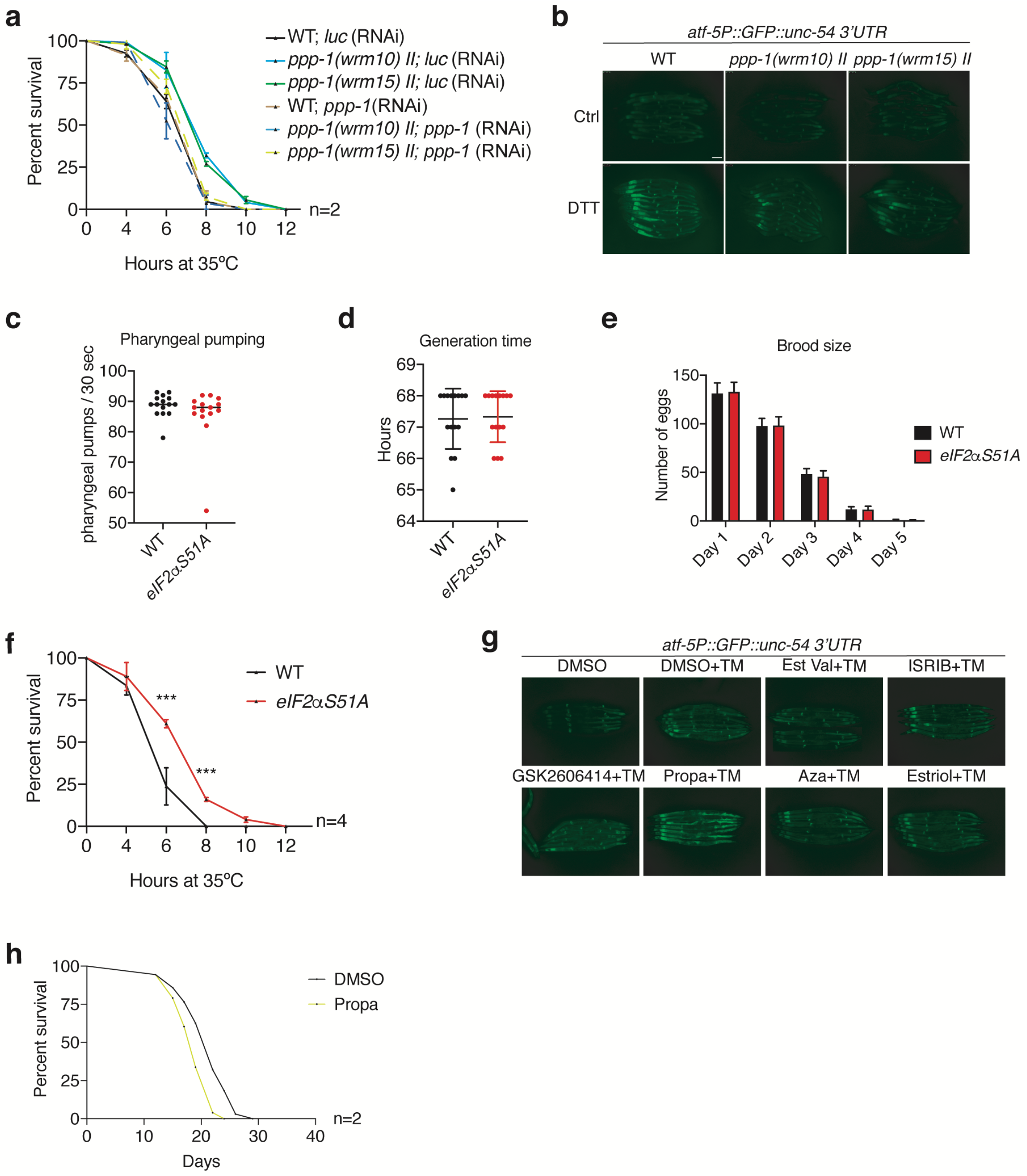


**Extended Data Fig. 2 | Characterization of Gcn(-) mutants and ISR modulator compounds.** **a**, Thermotolerance assays of day 1 WT and *ppp-1* mutants upon *ppp-1* RNAi treatment (error bars represent means ±SD, two-way ANOVA Dunnett's post hoc test; n=2). **b**, Fluorescence microscopy of day 1 WT and *ppp-1* mutants in the *atf-5P*::GFP::*unc-54* 3'UTR background, incubated without (Ctrl) or with 5 mM DTT for 2 h. Scale bar 75 µm. n=3. **c**, Pharyngeal pumping rates of day 1 WT and *eIF2αS51A* mutants (error bars represent means ±SD). **d**, Generation time of WT and *eIF2αS51A* mutants (error bars represent means ±SD). **e**, Brood size of WT and *eIF2αS51A* mutants (error bars represent means +SD). **f**, Thermotolerance assays of *eIF2αS51A*

mutants show significantly increased survival during heat stress compared to WT (error bars represent means ±SD, two-way ANOVA Sidak's post hoc test with ***$p<0.001$ versus WT controls; n=4). **g**, Fluorescence microscopy of WT *atf-5P*::GFP::*unc-54* 3'UTR reporter worms grown on NGM plates supplemented without or with 10 µM tunicamycin (TM) and the indicated compounds (20 µM) or 1% DMSO control only (Est Val = estradiol valerate, Propa = propafenone hydrochloride, Aza = azadirachtin). Scale bar 75 µm. **h**, Survival of WT worms grown on NGM plates supplemented with 1% DMSO or 20 µM propafenone hydrochloride (n=2). See Extended Data Table 1 for lifespan statistics. See Extended Data Table 2 for statistics on thermotolerance assays.

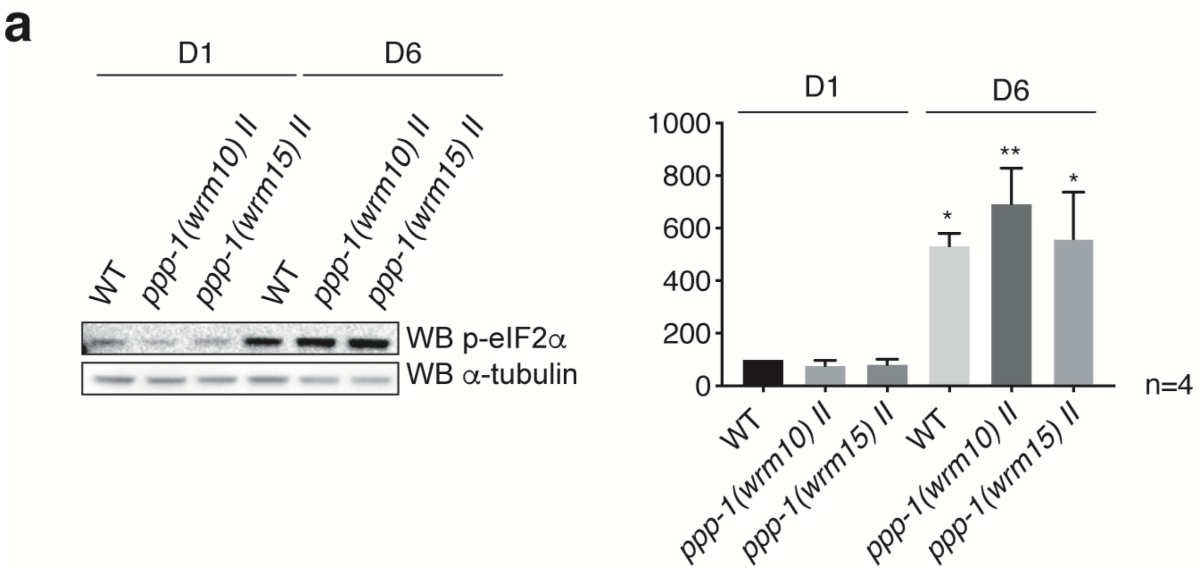


**Extended Data Fig. 3 | ISR induction in aged WT worms and *ppp-1* mutants. a**, Representative western blot of day 1 and day 6 WT and *ppp-1* mutants detecting phospho-eIF2α (Ser51) normalized to α-tubulin (error bars represent means +SEM, one-way ANOVA Tukey's post hoc test with *p<0.05 and **p<0.01 versus WT controls; n=4).

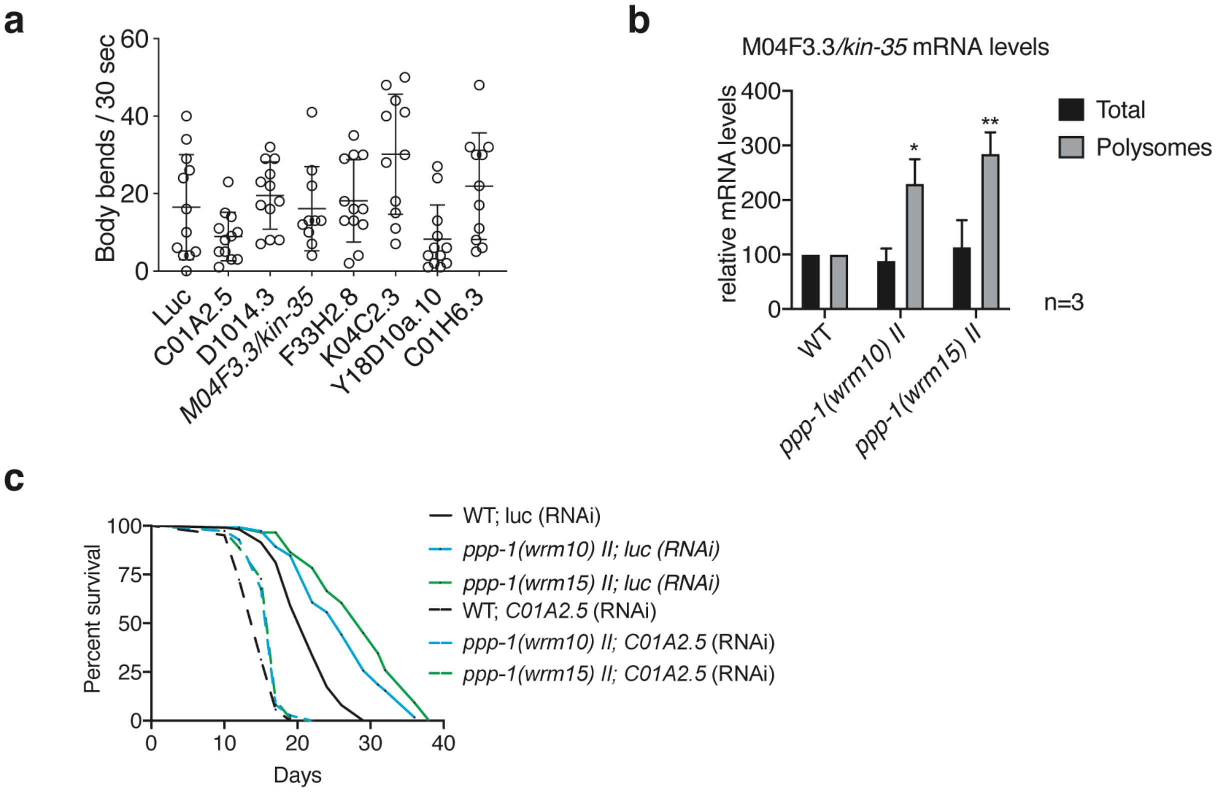


**Extended Data Fig. 4 | Supporting data for the RNAi screen for suppressors of *ppp-1* polyQ35 motility. a**, Control motility assays of day 6 WT polyQ35 worms after indicated RNAi treatments (error bars represent means ±SD, one-way ANOVA Dunnett's post hoc test. **b**, mRNA distribution of *kin-35* mRNA in total worm extracts and polysomes of day 1 WT and *ppp-1* animals (error bars represent means +SEM, two-way ANOVA Tukey's post hoc test with *p<0.05 and **p<0.01 versus WT controls; n=3) **c**, Survival of WT and *ppp-1* mutants upon RNAi knockdown of *C01A2.5* and control *luciferase*. See Extended Data Table 1 for lifespan statistics.

**Extended Data Table 1.** Lifespan statistics.

| Replicate | Strain | Treatment | Mean LS (days) | Difference (%) | dead/censored animals | P value | Reference control | P value | Reference control | Figure |
|---|---|---|---|---|---|---|---|---|---|---|
| 1 (11 02 18) | WT | | 21,42 | 0 | 132/32 | | | | | Fig. 1e |
| | *ppp-1(wrm10) II* | | 26,49 | 23 | 92/24 | p<0,0001 | vs WT | | | |
| | *ppp-1(wrm15) II* | | 26,9 | 25 | 111/43 | p<0,0001 | vs WT | | | |
| 2 (20/11/16) | WT | | 19,82 | 0 | 81/10 | | | | | |
| | *ppp-1(wrm10) II* | | 23,42 | 18 | 89/11 | p<0,0001 | vs WT | | | |
| 3 (10/10/16) | WT | | 20,36 | 0 | 94/9 | | | | | |
| | *ppp-1(wrm10) II* | | 25,09 | 23 | 75/25 | p<0,0001 | vs WT | | | |
| 2 (07 04 17) | WT | | 21,11 | 0 | 67/30 | | | | | |
| | *ppp-1(wrm15) II* | | 25,09 | 19 | 62/24 | p<0,0001 | vs WT | | | |
| 3 (29/03/17) | WT | | 21,26 | 0 | 121/14 | | | | | |
| | *ppp-1(wrm15) II* | | 24,6 | 16 | 113/7 | p<0,0001 | vs WT | | | |
| 1 (21 10 18) | WT | | 19,96 | 0 | 80/23 | | | | | |
| | *ppp-1(syb691) II* | | 22,75 | 14 | 83/31 | p<0,0001 | vs WT | | | |
| | *ppp-1(syb728) II* | | 23,04 | 15 | 73/37 | p<0,0001 | vs WT | | | |
| 2 (16 09 18) | WT | | 21,43 | 0 | 75/27 | | | | | Fig. 1f |
| | *ppp-1(syb691) II* | | 24,9 | 16 | 58/43 | p<0,0001 | vs WT | | | |
| | *ppp-1(syb728) II* | | 25,47 | 19 | 51/56 | p<0,0001 | vs WT | | | |
| 1 (28/08/16) | WT | | 22,11 | 0 | 73/9 | | | | | |
| | *gcn-2(wrm4) II* | | 24,28 | 10 | 73/11 | p = 0,010377 | vs WT | | | |
| | *pek-1(wrm7) X* | | 25,56 | 16 | 79/21 | p<0,0001 | vs WT | | | |
| 2(09 11 16) | WT | | 21,49 | 0 | 75/7 | | | | | |
| | *gcn-2(wrm4) II* | | 25,28 | 18 | 78/13 | p<0,0001 | vs WT | | | |
| | *pek-1(wrm7) X* | | 24,04 | 12 | 75/7 | p = 0,004266 | vs WT | | | |
| 3(30 10 16) | WT | | 20,03 | 0 | 71/23 | | | | | |
| | *gcn-2(wrm4) II* | | 24,13 | 20 | 69/9 | p = 0,007354 | vs WT | | | |
| | *pek-1(wrm7) X* | | 24,21 | 21 | 53/39 | p = 0,021492 | vs WT | | | |
| (19 01 19) | WT | | 19,79 | 0 | 83/33 | | | | | Fig. 1g |
| | *gcn-2(wrm4) II* | | 22,24 | 12 | 88/3 | p = 0,000218 | vs WT | | | |
| | *pek-1(wrm7) X* | | 22,8 | 15 | 89/32 | p<0,0001 | vs WT | | | |
| | *gcn-2(wrm4) II; pek-1(wrm7) X* | | 24,07 | 22 | 72/37 | p<0,0001 | vs WT | | | |
| 1 (28 06 17) | WT | luciferase | 20,31 | 0 | 49/15 | | | | | |
| | *ppp-1(wrm10) II* | luciferase | 25,87 | 27 | 44/27 | p<0,0001 | vs WT luc(RNAi) | | | |
| | *ppp-1(wrm15) II* | luciferase | 25,6 | 26 | 46/27 | p<0,0001 | vs WT luc(RNAi) | | | |
| | WT | *ppp-1* | 19,45 | -4 | 51/12 | p = 0,172333 | vs WT luc(RNAi) | | | |
| | *ppp-1(wrm10) II* | *ppp-1* | 20,37 | 0 | 63/4 | p = 0,957811 | vs WT luc(RNAi) | p<0,0001 | vs *ppp-1(wrm10) II; luc*(RNAi) | |
| | *ppp-1(wrm15) II* | *ppp-1* | 20,15 | -1 | 58/17 | p = 0,867597 | vs WT luc(RNAi) | p<0,0001 | vs *ppp-1(wrm15) II; luc*(RNAi) | |
| 2 (30 07 17) | WT | luciferase | 21,05 | 0 | 95/21 | | | | | Fig. 2a |
| | *ppp-1(wrm10) II* | luciferase | 26,28 | 25 | 65/33 | p<0,0001 | vs WT luc(RNAi) | | | |
| | *ppp-1(wrm15) II* | luciferase | 27,77 | 32 | 68/31 | p<0,0001 | vs WT luc(RNAi) | | | |
| | WT | *ppp-1* | 20,49 | -3 | 63/37 | p = 0,819891 | vs WT luc(RNAi) | | | |
| | *ppp-1(wrm10) II* | *ppp-1* | 21,48 | 2 | 72/24 | p = 0,044209 | vs WT luc(RNAi) | p<0,0001 | vs *ppp-1(wrm10) II; luc*(RNAi) | |
| | *ppp-1(wrm15) II* | *ppp-1* | 20,83 | -1 | 80/16 | p = 0,883359 | vs WT luc(RNAi) | p<0,0001 | vs *ppp-1(wrm15) II; luc*(RNAi) | |
| 1 (10 08 18) | WT | | 21,11 | 0 | 57/13 | | | | | Fig. 2b |
| | *ppp-1(wrm10) II / +* | | 26,23 | 24 | 42/32 | p<0,0001 | vs WT | | | |
| | *ppp-1(wrm15) II / +* | | 27,12 | 28 | 47/26 | p<0,0001 | vs WT | | | |
| 1 (09 06 19) | WT | | 19,57 | 0 | 95/14 | | | | | |
| | *eIF2a(syb1385) I* | | 22,18 | 13 | 102/20 | p<0,0001 | vs WT | | | |
| 2 (14 07 19) | *WT* | | 20,32 | 0 | 78/18 | | | | | Fig. 2e |
| | *eIF2a(syb1385) I* | | 24,22 | 19 | 75/25 | p<0,0001 | vs WT | | | |
| 3 (09 06 16) | WT | | 21,54 | 0 | 64/34 | | | | | |
| | *eIF2a(syb1385) I* | | 24,73 | 15 | 74/29 | p = 0,000142 | vs WT | | | |
| 4 (11 08 19) | WT | | 20,58 | 0 | 96/4 | | | | | |
| | *eIF2a(syb1385) I* | | 25,66 | 25 | 95/20 | p<0,0001 | vs WT | | | |
| 1 (6 11 18) | WT | DMSO | 20,53 | 0 | 67/39 | | | | | |
| | WT | Est Val D1 | 24,73 | 20 | 57/48 | p<0,0001 | vs DMSO | | | |
| | WT | Est Val D5 | 23,33 | 14 | 49/46 | p = 0,000948 | vs DMSO | | | |
| | WT | Est Val D10 | 22,36 | 9 | 55/46 | p = 0,009578 | vs DMSO | | | |
| 2 (27 06 18) | WT | DMSO | 21,06 | 0 | 70/20 | | | | | Fig. 2g |
| | WT | Est Val D1 | 24,62 | 17 | 53/33 | p<0,0001 | vs DMSO | | | |
| | WT | Est Val D5 | 24,06 | 14 | 60/33 | p<0,0001 | vs DMSO | | | |
| | WT | Est Val D10 | 24,26 | 15 | 54/38 | p = 0,000105 | vs DMSO | | | |
| 1 (6 11 18) | WT | DMSO | 20,53 | 0 | 67/39 | | | | | |
| | WT | Propa | 17,7 | -14 | 84/15 | p<0,0001 | vs DMSO | | | |
| 2 (27 06 18) | WT | DMSO | 21,06 | 0 | 70/20 | | | | | Extended Data Fig. 2h |
| | WT | Propa | 17,84 | -15 | 66/20 | p<0,0001 | vs DMSO | | | |
| 1 (16 06 19) | WT | luciferase | 22,76 | 0 | 60/41 | | | | | Fig. 4f |
| | *ppp-1(wrm10) II* | luciferase | 26,59 | 17 | 90/26 | p<0,0001 | vs WT RNAi(luc) | | | |
| | *ppp-1(wrm15) II* | luciferase | 26,87 | 18 | 68/29 | p<0,0001 | vs WT RNAi(luc) | | | |
| | WT | *M04F3.3* | 22,76 | 0 | 45/51 | p = 0,905931 | vs WT RNAi(luc) | | | |
| | *ppp-1(wrm10) II* | *M04F3.3* | 23,32 | 2 | 77/25 | p = 0,170039 | vs WT RNAi(luc) | p<0,0001 | vs *ppp-1(wrm10) II; luc*(RNAi) | |
| | *ppp-1(wrm15) II* | *M04F3.3* | 23,18 | 2 | 72/40 | p = 0,245433 | vs WT RNAi(luc) | p<0,0001 | vs *ppp-1(wrm15) II; luc*(RNAi) | |
| 2 (13 10 19) | WT | luciferase | 20,79 | 0 | 81/19 | | | | | |
| | *ppp-1(wrm10) II* | luciferase | 25,82 | 14 | 61/37 | p<0,0001 | vs WT RNAi(luc) | | | |
| | *ppp-1(wrm15) II* | luciferase | 26,61 | 28 | 63/43 | p<0,0001 | vs WT RNAi(luc) | | | |
| | WT | *M04F3.3* | 21,81 | 5 | 74/19 | p = 0,144674 | vs WT RNAi(luc) | | | |
| | *ppp-1(wrm10) II* | *M04F3.3* | 20,93 | 1 | 74/19 | p = 0,349183 | vs WT RNAi(luc) | p<0,0001 | vs *ppp-1(wrm10) II; luc*(RNAi) | |
| | *ppp-1(wrm15) II* | *M04F3.3* | 22,7 | 9 | 92/7 | p = 0,001631 | vs WT RNAi(luc) | p<0,0001 | vs *ppp-1(wrm15) II; luc*(RNAi) | |
| 1 (10 07 19) | WT | luciferase | 21,41 | 0 | 76/42 | | | | | Extended Data Fig. 4b |
| | *ppp-1(wrm10) II* | luciferase | 26,19 | 22 | 75/52 | p<0,0001 | vs WT RNAi(luc) | | | |
| | *ppp-1(wrm15) II* | luciferase | 28,59 | 34 | 62/46 | p<0,0001 | vs WT RNAi(luc) | | | |
| | WT | *C01A2.5* | 14,81 | -31 | 82/22 | p<0,0001 | vs WT RNAi(luc) | | | |
| | *ppp-1(wrm10) II* | *C01A2.5* | 16,33 | -24 | 83/36 | p<0,0001 | vs WT RNAi(luc) | | | |
| | *ppp-1(wrm15) II* | *C01A2.5* | 16,31 | -24 | 84/26 | p<0,0001 | vs WT RNAi(luc) | | | |
| 1 (25 08 19) | WT | luciferase | 21,62 | 0 | 78/29 | | | | | Fig. 4g |
| | *eIF2$\alpha$ (syb1385) I* | luciferase | 24,87 | 15 | 77/21 | p<0,0001 | vs WT RNAi(luc) | | | |
| | WT | *M04F3.3* | 21,95 | 2 | 79/23 | p = 0,988349 | vs WT RNAi(luc) | | | |
| | *eIF2$\alpha$ (syb1385) I* | *M04F3.3* | 22,97 | 6 | 79/35 | p = 0,066634 | vs WT RNAi(luc) | p = 0,002449 | vs eIF2$\alpha$(syb1385) I; *luc*(RNAi) | |
| 2 (12 12 19) | WT | luciferase | 18,98 | 0 | 90/13 | | | | | |
| | *eIF2$\alpha$ (syb1385) I* | luciferase | 23,6 | 24 | 68/39 | p<0,0001 | vs WT RNAi(luc) | | | |
| | WT | *M04F3.3* | 18,8 | -1 | 85/25 | p = 0,930616 | vs WT RNAi(luc) | | | |
| | *eIF2$\alpha$ (syb1385) I* | *M04F3.3* | 21,25 | 12 | 78/26 | p = 0,001145 | vs WT RNAi(luc) | p = 0,001458 | vs eIF2$\alpha$(syb1385) I; *luc*(RNAi) | |
| 3 (14 10 19) | WT | luciferase | 20,4 | 0 | 66/18 | | | | | |
| | *eIF2$\alpha$ (syb1385) I* | luciferase | 23,87 | 17 | 69/24 | p<0,0001 | vs WT RNAi(luc) | | | |
| | WT | *M04F3.3* | 18,88 | -7 | 67/27 | p = 0,039434 | vs WT RNAi(luc) | | | |
| | *eIF2$\alpha$ (syb1385) I* | *M04F3.3* | 19,32 | -5 | 59/25 | p = 0,111907 | vs WT RNAi(luc) | p<0,0001 | vs eIF2$\alpha$(syb1385) I; *luc*(RNAi) | |
| 4 (17 11 19) | WT | luciferase | 21,12 | 0 | 68/31 | | | | | |
| | *eIF2$\alpha$ (syb1385) I* | luciferase | 24,24 | 15 | 47/17 | p = 0,000149 | vs WT RNAi(luc) | | | |
| | WT | *M04F3.3* | 21,66 | 3 | 61/23 | p = 0,386793 | vs WT RNAi(luc) | | | |
| | *eIF2$\alpha$ (syb1385) I* | *M04F3.3* | 21,64 | 2 | 63/21 | p = 0,556019 | vs WT RNAi(luc) | p = 0,000285 | vs eIF2$\alpha$(syb1385) I; *luc*(RNAi) | |

**Extended Data Table 2.** Thermotolerance statistics.

| Time (Hours) | WT | | | | *ppp-1(wrm10)* II | | | | *ppp-1(wrm15) II* | | | |
|---|---|---|---|---|---|---|---|---|---|---|---|---|
| **0** | 100 | 100 | 100 | 100 | 100 | 100 | 100 | 100 | 100 | 100 | 100 | 100 |
| **4** | 83 | 95 | 88 | 100 | 98 | 97 | 100 | 98 | 96 | 100 | 100 | 100 |
| **6** | 50 | 53 | 40 | 32 | 86 | 77 | 94 | 70 | 82 | 68 | 87 | 72 |
| **8** | 0 | 0 | 0 | 0 | 22 | 14 | 30 | 24 | 36 | 15 | 49 | 24 |
| **10** | 0 | 0 | 0 | 0 | 0 | 0 | 0 | 8 | 0 | 0 | 0 | 4 |
| **12** | 0 | 0 | 0 | 0 | 0 | 0 | 0 | 0 | 0 | 0 | 0 | 0 |
| | Fig. 1H | | | | Fig. 1H | | | | Fig. 1H | | | |

| | Luciferase (RNAi) | | | | | | ppp-1 (RNAi) | | | | | |
|---|---|---|---|---|---|---|---|---|---|---|---|---|
| **Time (Hours)** | **WT** | | ***ppp-1(wrm10)* II** | | ***ppp-1(wrm15) II*** | | **WT** | | ***ppp-1(wrm10)* II** | | ***ppp-1(wrm15) II*** | |
| **0** | 100 | 100 | 100 | 100 | 100 | 100 | 100 | 100 | 100 | 100 | 100 | 100 |
| **4** | 91 | 94 | 100 | 98 | 100 | 96 | 89 | 94 | 98 | 100 | 96 | 100 |
| **6** | 67 | 61 | 75 | 90 | 87 | 82 | 67 | 66 | 45 | 60 | 70 | 76 |
| **8** | 9 | 0 | 33 | 31 | 26 | 28 | 9 | 0 | 7 | 0 | 9 | 6 |
| **10** | 0 | 0 | 4 | 4 | 7 | 4 | 0 | 0 | 0 | 0 | 0 | 0 |
| **12** | | 0 | 0 | 0 | 0 | 0 | 0 | 0 | 0 | 0 | 0 | 0 |

Data depicted in yellow belong to Extended Data Figure 1B

| Time (Hours) | WT | | | | *eIF2α (syb1385) I* | | | |
|---|---|---|---|---|---|---|---|---|
| **0** | 100 | 100 | 100 | 100 | 100 | 100 | 100 | 100 |
| **4** | 77 | 88 | 88 | 81 | 81 | 83 | 94 | 98 |
| **6** | 15 | 15 | 38 | 27 | 60 | 63 | 63 | 58 |
| **8** | 0 | 0 | 0 | 0 | 15 | 15 | 17 | 17 |
| **10** | 0 | 0 | 0 | 0 | 4 | 6 | 2 | 4 |
| **12** | 0 | 0 | 0 | 0 | 0 | 0 | 0 | 0 |
| | Fig. 2G | | | | Fig. 2G | | | |

**Extended Data Table 3.** List of polysome-associated mRNAs in WT and *ppp-1* mutants. Data displayed as fold change in WT versus *ppp-1.*

Please see excel file uploaded as Auxiliary Supplementary Information.

**Extended Data Table 4.** Worm strains used in this study.

| Strain name | Genotype | Backgrossed to N2 | Source |
|---|---|---|---|
| MSD331 | Bristol N2 | | CGC |
| MSD283 | *ppp-1(wrm10) II* | 4x | this study |
| MSD310 | *ppp-1(wrm15) II* | 4x | this study |
| MSD275 | *gcn-2(wrm4) II* | 4x | this study |
| MSD302 | *pek-1(wrm7) X* | 4x | this study |
| MSD314 | ldIs(atf-5P::GFP::unc-54 3'UTR) | 2x | Blackwell lab |
| MSD317 | *ldIs(atf-5P::GFP::unc-54 3'UTR); ppp-1(wrm10) II* | 2x | this study |
| MSD456 | *ldIs(atf-5P::GFP::unc-54 3'UTR); ppp-1(wrm15) II* | 2x | this study |
| SYB728 | *ppp-1(syb728) II* CR-L216F (wrm15) | n/a | Suny Biotech |
| SYB691 | *ppp-1(syb691) II* CR-N295I (wrm10) | n/a | Suny Biotech |
| MSD406 | *rsks-1(sv31) III* | 2x | Hubbard lab |
| MSD441 | *mIs133[unc-54p::Q35:YFP]* | 2x | CGC |
| MSD442 | *mIs133[unc-54p::Q35:YFP]; ppp-1(wrm10)* | 2x | this study |
| MSD457 | *mIs133[unc-54p::Q35:YFP]; ppp-1(wrm15)* | 2x | this study |
| MSD412 | *gcn-2(ok871) II; pek-1(ok275) X* | 2x | this study |
| AA4685 | *ifg-1(cxTi9279)* | n/a | Antebi Lab |
| SYB1385 | *eIF2α(syb1385) I* | 2x | Suny Biotech |
| CF512 | *fer15(b26) II; fem-1(hc17) IV* | n/a | CGC |

**Extended Data Table 5.** Genotyping methods and primers used in this study.

| Genotype | Genotyping Method | Primer sequence |
|---|---|---|
| *ppp-1(wrm10) II*<br>*ppp-1(wrm15) II*<br>*gcn-2(wrm4) II*<br>*pek-1(wrm7) X* | TaqMan SNP mapping | Manufactured by Applied Biosysems |
| *ppp-1(syb691) II* CR-N295I (wrm10) | PCR | For TCCTTGTCAATCTGAATGGA<br>Rev CCATCGATCAAATTAACTTCA |
| *ppp-1(syb728) II* CR-L216F (wrm15) | PCR | For GCAGTTACAAATCCATTTT<br>Rev CCTTAATAATTCTAATTTTTACA |
| *gcn-2(ok871) II* | PCR | For GCATGACATGGCAATGATTCATCG<br>Rev GTACTTTGCAATGATTTTGAGGCG |
| *pek-1(ok275) X* | PCR | For CTGAGAAGGCAACGCTCTCT<br>Rev ATCACCGCTACTCTGGATGG |
| *eIF2α(syb1385) I* | PCR | For ATTGGTAGTTTATCTCATTTAAT<br>Rev TTCTCCTTAATATCTGCACT |
